# The genomic basis of host and vector specificity in non-pathogenic trypanosomatids

**DOI:** 10.1101/2022.01.05.475049

**Authors:** Guy Oldrieve, Beatrice Malacart, Javier López-Vidal, Keith Matthews

## Abstract

The ability of trypanosome parasites to survive and sustain infections is dependent on diverse and intricate immune evasion mechanisms. Pathogenic trypanosomes often have broad host niches that preclude identification of host specific adaptations. In contrast, some non-pathogenic species of the genus *Trypanosoma* have highly specific hosts and vectors. *Trypanosoma theileri*, a non-pathogenic parasite of bovines, has a predicted surface protein architecture that likely aids survival in its mammalian host, distinct from the dominant variant surface glycoprotein coat of pathogenic African trypanosomes. In both species, their surface proteins are encoded by genes which account for ∼10% of their genome. A non-pathogenic parasite of sheep, *Trypanosoma melophagium*, is transmitted by the sheep ked and is closely related to *T. theileri*. To explore host and vector specificity between these closely related species, we sequenced the *T. melophagium* genome and transcriptome and an annotated draft genome was assembled. *T. melophagium* was compared to 43 kinetoplastid genomes, including *T. theileri. T. melophagium* and *T. theileri* have an AT biased genome, the greatest bias of publicly available trypanosomatids. This trend may result from selection acting to decrease the genome nucleotide cost. The *T. melophagium* genome is 6.3Mb smaller than *T. theileri* and large families of proteins, characteristic of the predicted surface of *T. theileri,* were found to be absent or greatly reduced in *T. melophagium.* Instead, *T. melophagium* has modestly expanded protein families associated with the avoidance of complement-mediated lysis. The genome of *T. melophagium* contains core genes required for development, glycolysis, RNA interference, and meiotic exchange, each being shared with *T. theileri*. Comparisons between *T. melophagium* and *T. theileri* provide insight into the specific adaptations of these related trypanosomatids to their distinct mammalian hosts and arthropod vectors.

**Author summary:** Non-pathogenic trypanosomes can have narrow host niches, with closely related trypanosome species expanding into distinct mammalian host and insect vectors. *T. theileri*, a non-pathogenic trypanosome of bovines, is predicted to have an intricate cell surface which allows it to evade the immune response of its mammalian host. In contrast, *T. melophagium* is closely related to *T. theileri* but infects sheep and is transmitted by the sheep ked rather than tabanid flies that transmit *T. theileri*. Here, we sequence and assemble the *T. melophagium* genome to identify the genomic basis of host and vector specificity in these non-pathogenic trypanosomes. We confirm the two species are closely related, however, *T. melophagium* has a smaller genome than *T. theileri*. Most of the discrepancy in genome size is due to an expansion of putative cell surface genes in *T. theileri*. The differential investment in cell surface proteins could be due to a focus on adaptation to the mammalian host in *T. theileri* and the insect host in *T. melophagium*.

**Data summary:** The genomes, transcriptomes and proteomes used in this study were accessed from the TriTrypDB repository or NCBI. *T. theileri* genome sequencing data was downloaded from NCBI SRA (SRR13482812). *T. melophagium* data generated during this study is available from the NCBI BioProject PRJNA786535.

**Repositories:** *T. melophagium* DNA and RNA sequencing data, along with the draft genome assembly and its annotation, can be found under the NCBI BioProject PRJNA786535.

## 5. Introduction

Trypanosomatidae are a family of single celled eukaryotes, characterised by a specialised mitochondrial genome, the kinetoplast. Trypanosomatidae are monoxenous (single host) or dixenous (two host) species. Dixenous trypanosomatids are obligate parasites of a broad diversity of animals and plants whilst monoxenous species are largely restricted to insects (1). However, the taxonomy of trypanosomatids cannot be distilled into these two broad categories, as many monoxenous species opportunistically infect vertebrates (2) and some dixenous species have subsequently reverted to a monoxenous lifecycle (3).

Expansion into vertebrate hosts gave rise to clades of trypanosomatids which represent medical and veterinary threats. Notably, *Trypanosoma cruzi* and *Leishmania* spp. cause important diseases in humans, Chagas disease and Leishmaniasis, respectively. Also, *Trypanosoma brucei* has been subjected to intense molecular and cytological study as two of its subspecies, *Trypanosoma brucei gambiense* and *Trypanosoma brucei rhodesiense*, are causative agents of Human African Trypanosomiases (HAT) (4). In addition to the medical impact of human African trypanosomes, other African trypanosome species such as *T. b. brucei*, *Trypanosoma vivax* and *Trypanosoma congolense* cause morbidity and mortality in livestock, constraining agricultural development (5).

Trypanosomatids diverged from other eukaryotes over a billion years ago (6, 7) and possess unique adaptations at the genome level (8). The *T. brucei* genome comprises 11 megabase chromosomes (9) along with ∼5 intermediate chromosomes and ∼100 mini-chromosomes (10, 11). Nuclear DNA is highly compact, and genes are organised into polycistronic units, the primary transcripts of which are trans-spliced and polyadenylated to resolve mature mRNA (12, 13). Their distinct mitochondrial genome, the kinetoplast, consists of mini (14, 15) and maxicircles (16, 17) that rely upon RNA editing to generate functional mRNA (8).

The divergence in trypanosomatid morphology, transmission, and lifecycle development has facilitated an adaption to a diverse range of hosts and vectors. The different surface protein environments of *T. brucei*, *T. cruzi* and *Leishmania* spp. exemplify the variant strategies adopted by these parasites to evade the immune systems of their hosts and vectors. For example, *T. brucei* is extracellular and proliferates in the blood and tissue of mammals. Its cell surface is encoded by a very extensive repertoire of variant surface glycoproteins (VSGs) (10, 18). VSG variation is essential to sustain long-term infections, during which antigenically distinct VSG protein types dominate at each peak, facilitating host immune evasion (19, 20). *T. cruzi* is an intracellular parasite of wild and domestic mammals. The nuclear genome contains an expanded family of mucin genes that represent up to 6% of the genome (21) and, along with trans-sialidases (22), these genes enable sustained infections. *Leishmania* spp. are intracellular trypanosomatids whose cell surface is covered by a thick layer of glycoconjugates, including families of major surface proteases (MSPs) (23).

Studies have largely focused on pathogenic dixenous trypanosomatids, which can hold broad host niches, capable of infecting multiple mammalian species (24). In contrast, non-pathogenic dixenous trypanosomes can be highly specific to their host and vector. These species represent an attractive model to study the basis of host and vector specificity.

*Trypanosoma theileri* is a non-pathogenic bovine parasite which has a reported prevalence of 80% in cattle in the US and Europe when screened via culture-based methods (25–28). *T. theileri* is transmitted by tabanid flies (29). It can cause lifelong infections but remains at an extremely low parasitaemia in immunocompetent animals, indicating the presence of an effective host immune evasion mechanism or strict self-imposed population control, which prevents overt disease (30, 31). Experimentally, *T. theileri* can sustain an infection for at least 12 weeks (28) which, combined with their non-pathogenic nature, has stimulated development of *T. theileri* as a potential vaccine delivery vehicle (28). The genome of *T. theileri* encodes five predicted surface protein families. These are four unique protein families, *T. theileri* putative surface proteins (TTPSPs) and one MSP family (32). Together these genes represent 9% of the genome of *T. theileri*, comparable to the representation of the VSG gene family in *T. brucei.* Lastly, the trans-sialidases that characterise *T. cruzi* were also found to be highly expressed in *T. theileri*. These findings led to the suggestion of a novel immune evasion mechanism in *T. theileri,* contrasting with the well-known system in African trypanosomes and distinct from that in *T. cruzi* or *Leishmania* (32).

*Trypanosoma melophagium* is another non-pathogenic trypanosome of the subgenus *megatrypanum*, transmitted between sheep via the sheep ked. This flightless insect vector has been eradicated from much of its original geographic distribution due to widespread pesticide use. However, where the sheep ked persists, it often carries *T. melophagium* (33, 34). In a study of organic sheep farms, *T. melophagium* was found to be present in 86% of keds, however, blood smears from sheep on the same farms did not detect trypanosomes (34). Other surveys via blood culture found 7.8% of sheep to be infected with *T. melophagium* (35). It has historically been argued that *T. melophagium* is a monoxenous parasite of the sheep ked and that the mammalian host is obsolete for transmission (36–38). However, extensive studies demonstrated a mammalian host is required (39). Experimental infections of sheep with *T. melophagium* suggested the longest infection lasts three months and there is no lasting immunity as sheep can be reinfected with *T. melophagium* after several months of isolation (33, 39, 40)

Molecular markers place *T. melophagium* as a close relative to *T. theileri* (34). SSU rRNA is ∼98% similar between *T. theileri* and *T. melophagium* isolates (33). Presumably the divergence of *T. theileri* and *T. melophagium* is associated with their discrete host niches (33, 34). Nonetheless, *T. theileri* and *T. melophagium* undergo a similar transmission cycle where metacyclic forms are produced in the insect hindgut and the infective forms are then believed to be transmitted to their mammalian host via the mouth, by ingestion of insect faeces or the whole insect body. Trypanosomes then invade their mammalian hosts and proliferate in the blood, before being taken up as a bloodmeal by their insect vector (29, 39).

A notable contrast in the biology of these parasites is the divergence in the life history of their vectors. Sheep keds, which transmit *T. melophagium*, spend their entire life attached to either the skin or wool and hair of sheep. Both male and female keds feed on their mammalian host (41). Tabanids, which transmit *T. theileri*, breed and lay their eggs in soil, water, or trees. The larvae and pupae stages live on vegetation and soil. Only female adults feed on mammalian blood, which is essential for egg production. Although adult tabanids show considerable adaptation to blood feeding, they also feed on the sugars of plants (42).

Here we derive the *T. melophagium* genome and compare its features with *T. theileri* to provide insight into the biological specificity exhibited by each parasite in the context of their close phylogenetic relationship.

### 6.1. Methods

#### Trypanosome culture, DNA/RNA extraction and sequencing

*T. melophagium* was isolated from sheep blood collected on the island of St Kilda, Scotland (Kindly provided by Professor Josephine Pemberton, University of Edinburgh). Whole blood was collected into heparinized vacutainers and used within 2 days. 1 ml of blood was diluted with 5 volumes of a 50% mix of HMI9 supplemented with 20% FBS and MDBK conditioned medium. All cultures were kept at 37°C and were examined microscopically every 3 days for 6 weeks. After propagation of *T. melophagium* by culturing of the blood sample, the specimens were transferred and co-cultured with fibroblast-like primary cells as feeder cells, isolated from the same blood sample. Due to the short lifespan of these primary cultured cells, *T. melophagium* was subsequently co-cultivated with MDBK cells and then progressively adapted to axenic conditions with a 50% mix of HMI9 and MDBK conditioned medium.

DNA was extracted from cultured *T. melophagium* using a MagAttract high molecular weight DNA kit, following the manufacturer’s instructions (Qiagen) and cleaned via ethanol precipitation. The DNA was sequenced with Oxford Nanopore Technology’s (ONT) MinION (R9.4.1), following the Nanopore Rapid Sequencing protocol. Base-calling was performed in high accuracy mode using Guppy (available at https://community.nanoporetech.com/) which produced 1.059 gigabases (Gb) of data. PycoQC was used to visualise the data (43). The same DNA was sequenced with BGI’s DNBseq (4.201Gb, 150 base pair (bp) reads). RNA was extracted with the RNeasy mini kit (Qiagen) including a DNAse step, following the manufacturer’s instructions and sequenced with BGI’s DNBseq (5.019Gb, 100bp reads). Raw DNA and RNA DNBseq reads were trimmed with Trimmomatic (44).

*T. theileri* sequencing data was downloaded from NCBI. 170bp genomic (SRR13482812) were used.

#### *T. melophagium* genome assembly and annotation

Jellyfish and GenomeScope were used to provide a k-mer based estimate of the genome size and heterozygosity using the short DNA reads described above (45, 46).

Nanopore long reads were assembled using Wtdbg2 (47). For the polishing steps, BWA-MEM (48) was used to align short reads and Minimap2 (49) was used to align ONT reads. ONT reads were aligned to the Wtdbg2 draft assembly and three iterations of Racon (50) followed by one round of Medaka (available at https://github.com/nanoporetech/medaka) were performed. DNBseq reads were mapped to the Medaka polished assembly, and two iterations of Racon were performed. Short and long reads were aligned to the Racon polished assembly to complete two final iterations of polishing with Pilon (51). At each stage of polishing, the draft genome was assessed using scaffold_stats.pl (available at https://github.com/sujaikumar/assemblage) and BUSCO (52).

Both sets of reads were mapped to the draft assembly. Each contig was subjected to a DIAMOND blastx search against the InterProScan database (53, 54). The resulting alignments and DIAMOND hits were visualised with BlobTools (55). Two contigs were outliers in comparison to the rest of the assembly at under 100x coverage and consisting of only 3,975 and 1,892 base pairs (Fig. S2a). These contigs were removed from the assembly. BUSCO and BlobTools were re-run on this trimmed assembly to assess the final assembly’s completeness and coverage (Supplementary Fig. S2b).

Repeat sequences in the genome were identified and soft-masked using RepeatModeler2 and RepeatMasker (56). BRAKER2 was used to annotate the soft-masked genome in ETP mode. The OrthoDB v10 protozoa database was utilised for protein hints (57) with the addition of the *T. theileri* proteome and RNAseq evidence (53, 58–68). BRAKER2 produced the protein and transcript files utilised in the following analysis. The *T. melophagium* proteome was functionally annotated using InterProScan (54) using the Pfam and SignalP databases (69, 70). Genome conservation of collinearity was compared using D-Genies (71).

Publicly available genomes, transcriptomes and proteomes were accessed from TriTrypDB (72) along with the *Phytomonas* EM1 assembly which was accessed from NCBI (73). The quality of the proteomes were assessed using BUSCO; only isolates which had > 85% complete proteomes were included, which left 43 isolates along with *T. melophagium*. A list of all the isolates used in this study can be found in the Supplementary File S1. The assembly statistics of the 44 genomes were assessed using scaffold_stats (available at https://github.com/sujaikumar/assemblage).

The genomes from each of these trypanosomatid isolates were screened for transfer RNA genes using tRNAscan-SE (74). The outputs of these results were used to infer the strength of selection acting on translational efficiency and nucleotide cost for each isolate, along with the background mutation bias, using CodonMuSe (75, 76). Each isolate was assessed using 1) every CDS, 2) single copy universal orthologs (identified in the orthologous protein clustering steps below) (n=992) and 3) single copy universal orthologs which are essential for every stage of the *T. brucei* life cycle (n=158). The last list of genes were identified by screening the universal single copy orthologs for genes which had a significant reduction in transcript levels in every library of an RNAi phenotype screen (>1.5 log fold decrease) (77).

#### Orthology inference

Orthologous proteins from 44 trypanosomatid proteomes were identified with OrthoFinder (78) and protein clusters were summarised with KinFin (79), using InterProScan annotations based on the Pfam and signalP databases (54, 69, 70). A minimal cut-off threshold was not applied to the orthogroup annotation. The orthogroup annotation summary can be found in Supplementary File S2. A species tree was produced by STAG and STRIDE, as part of the OrthoFinder analysis (80, 81) which was visualised with iTOL (82). STRIDE identified *Bodo saltans* as the best root for the consensus species tree.

To confirm the absence of VSGs in *T. melophagium*, all CDS sequences labelled as ‘VSG’ in the *T. brucei* TREU927/4 reference genome were downloaded from TriTrypDB. A blastn search was performed with the *T. melophagium* genome as the database using a loose cut-off (e-value = 1e-5). A similar search was performed to confirm the reduced TTPSP counts in *T. melophagium*. For this, the *T. theileri* transcripts were used to query a database made from the *T. melophagium* transcripts (e-value = 1e-25).

The cell surface orthogroups in this study were annotated with the orthogroups from Kelly *et al*. 2017 (32) based on the orthogroup membership of genes in the two analyses. Unless stated otherwise, all of the figures in this study were plotted in R (83) using ggplot2 (84) and ggrepel (85).

The full list of tools used in this study, and the options used to run those tools, can be found in Table S1.

### 6.2. Results

#### *T. melophagium* genome assembly

An initial assessment of the *T. melophagium* genome, via k-mer counting, predicted that it was smaller than that of *T. theileri* (22.3 Mb and 27.6 Mb, respectively), this variation being observed in its repeat and unique sequence (Table 1). Notably, both genomes are predicted to have extremely low heterozygosity (0.30 and 0.41 for *T. melophagium* and *T. theileri,* respectively) in comparison to other *Trypanosoma* isolates (86). Large gene families, such as the TTPSPs, only account for ∼10% of the *T. theileri* genome and so are predicted to have a minor effect in the heterozygosity calculation. The k-mer counting prediction was similar in size to the final assembly (Table 1). The *T. melophagium* assembly consisted of 64 contigs in comparison to 253 for *T. theileri*. BUSCO assessments suggest both assemblies are 100% complete, although *T. theileri* is slightly fragmented (Table 1).

**Table 1:** Genome assessment. The assemblies used in this assessment represent the final draft of the *T. melophagium* assembly produced during this study and the *T. theileri* assembly available from TriTrypDB (32, 72). The k-mer spectra plots associated with the first section of the table are found in Fig. S1.

|  | *T. melophagium* | *T. theileri* (32) |
| --- | --- | --- |
| **k-mer based genome survey** |  |  |
| Heterozygosity (%) | 0.30 | 0.41 |
| Genome / repeat /<br>unique length (Mbp) | 22.28 / 4.16 / 18.13 | 27.62 / 7.97 / 19.65 |
| **Genome assembly statistics (for scaffolds longer than 200 bp)** |  |  |
| Number of contigs | 64 | 253 |
| Span (Mbp) | 23.3 | 29.6 |
| Min / Max / Mean<br>(bp) | 5,186 / 1,230,212 / 364,120 | 791 / 1,635,300 / 117,005 |
| N50 | 505,851 | 517,122 |
| GC (%) | 41.2 | 40.1 |
| **Genome BUSCO assessment** |  |  |
| Complete/ Single<br>copy/ Duplicated/<br>Fragmented | 100 / 100 / 0 / 0 | 99.2 / 99.2 / 0 / 0.8 |
| **Annotation** |  |  |
| Number of genes | 10,057 | 11,312 |
| **Annotation BUSCO assessment** |  |  |
| Complete/ Single<br>copy/ Duplicated/<br>Fragmented | 100 / 100 / 0 / 0 | 99.2 / 99.2 / 0 / 0.8 |

The *T. melophagium* and *T. theileri* genomes were aligned to highlight the conservation of collinearity. The conservation is more similar to the conservation between the species *T. brucei* TREU927/4 *and T. congolense* IL3000 2019 than between isolates of the same species, *T. brucei* TREU927/4 and *T. brucei* Lister 427 (Fig. 1 & Fig. S3). The *T. melophagium* genome was annotated with 10,057 proteins, in comparison to 11,312 in *T. theileri* (Fig. 2c & Table 1) (32).

**Figure 1:**
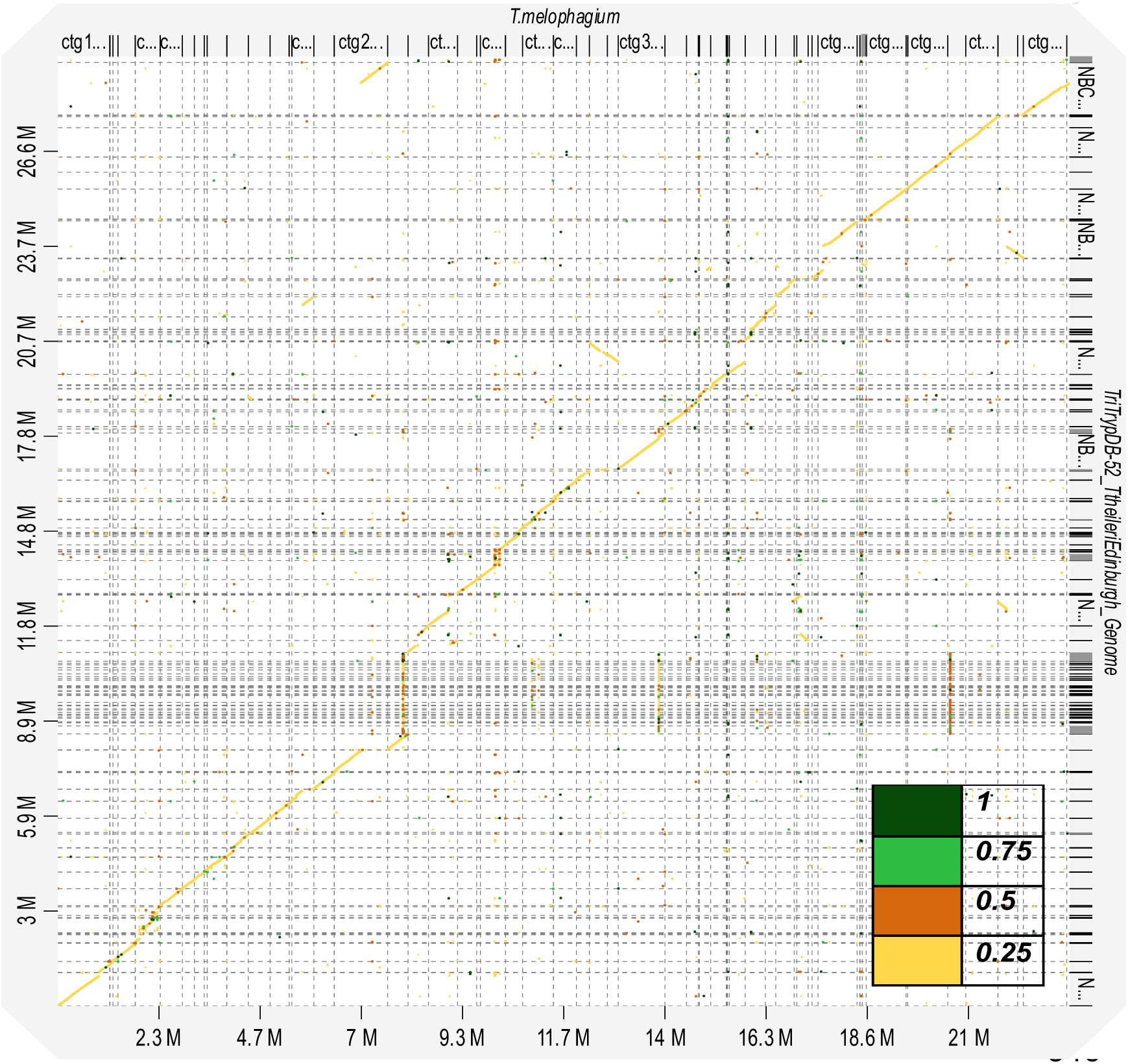
Synteny of the *T. melophagium* and *T. theileri* genome sequences highlights conservation and identity between the two species. The legend refers to the percentage identity between the sequences.

**Figure 2:**
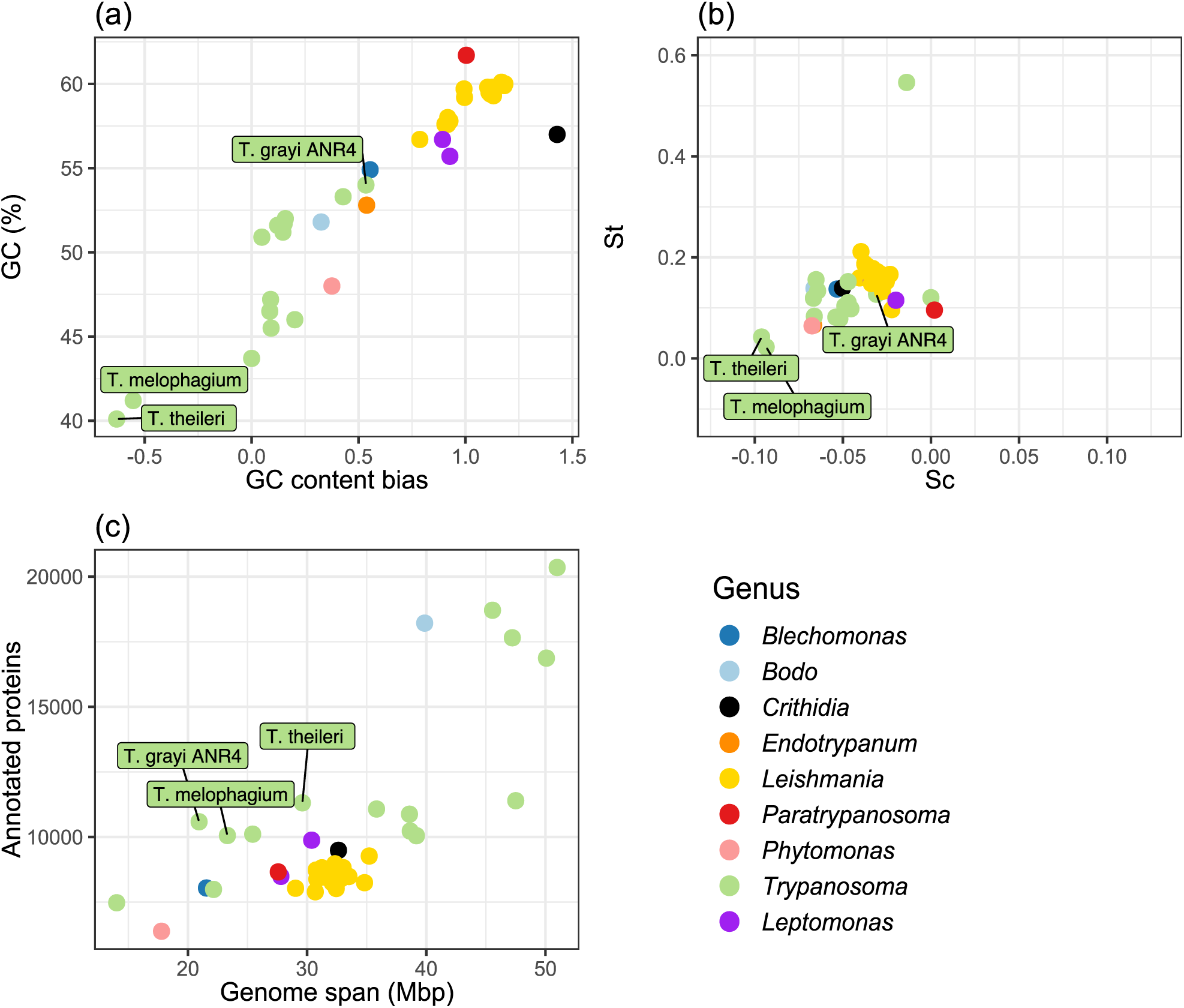
(a) GC content across the whole genome and GC content bias in the CDS of trypanosomatid universal single copy orthologues (n=992). GC content (GC) > 0 = GC content bias. GC < 0 = AT content bias. (b) Selection acting on translational efficiency (St) and selection acting on nucleotide cost (Sc) in trypanosomatid universal single copy orthologues. Sc > 0 = Selection acting to increase codon nucleotide cost. Sc < 0 = Selection is acting to decrease codon nucleotide cost. St > 0 = Selection is acting to increase codon translational efficiency. St < 0 = Selection acting to decrease codon translational efficiency. (c) Counts of annotated protein sequences of publicly available trypanosomatids compared by genome size.

Trypanosomatid genomes, proteomes and transcriptomes were downloaded from TriTrypDB (72). Proteome completeness was assessed with BUSCO. Those above 85% complete were retained, leaving 44 isolates from 9 genus (Supplementary File S1). *T. theileri* and *T. melophagium* have some of the smallest genomes in this study group and have the lowest GC content (40.1% and 41.2%, respectively), contrasting with the trypanosomatid mean of 48.4% (Fig. 2a).

Environmental temperature, generation time, neutral drift, gene splicing, tRNAs, translational accuracy/efficiency and protein folding have all been hypothesised to cause codon bias. However, codon bias can be influenced by nutrient availability (76). Species with low nitrogen availability, such as the plant trypanosomatid, *Phytomonas*, have an AT rich genome, potentially to mitigate the lack of nitrogen in their plant hosts (76). Selection acting on genome nucleotide cost (Sc) is in competition with selection acting to alter the translational efficiency of the genome (St) (75, 76). Based on universal single copy orthologues, selection pressure acting on the translational efficiency is minimal for both *T. theileri* and *T. melophagium*. In contrast, they show the greatest predicted selection pressure acting to reduce the nucleotide cost of any trypanosomatid genome. The AT biased content can be interpreted as a remodelling of the *T. theileri* and *T. melophagium* genomes to reduce the cost of the genome (Fig. 2 a & b). This pattern held true when every CDS was compared (Fig. S4 a & b) and in universal single copy orthologs which are essential for every life cycle stage of *T. brucei* (Fig. S4 c & d).

#### Orthologous protein clustering and phylogenetic inference

Orthologous clustering identified genes that descended from a gene in the last common ancestor of the 44 trypanosomatid proteomes used in this study (Fig. 2, Supplementary File S1). From the 44 proteomes, 18,274 orthogroups were identified (96.5% of the proteins used in this study were included in one of these orthogroups), 992 orthogroups were single copy and contained all isolates.

A species tree was generated as part of the orthologous protein clustering. Using genetic markers, previous studies have noted the similarity between *T. melophagium* and *T. theileri* (33, 34). Based on 2,312 gene trees, *T. theileri* is the closest isolate to *T. melophagium* and groups with the stercorarian trypanosomes, which include *T. cruzi*, *Trypanosoma grayi* and *Trypanosoma rangeli,* rather than with salivarian trypanosomes such as *T. brucei* (Fig. 3). *T. grayi* is the closest isolate to the *T. melophagium* and *T. theileri* clade and is closer in size to the genome span of *T. melophagium* than to *T. theileri* (Fig. 2c). *T. melophagium* and *T. theileri* do seem to be closely related species with relatively short branch lengths (0.093 and 0.088 substitutions per site, respectively). In comparison, *T. congolense* is more divergent from *T. brucei* (0.249) whilst the isolates within *T. brucei* (*T. brucei brucei* Lister 427:0.0009, *T. brucei evansi* STIB805: 0.003, *T. brucei brucei* TREU927/4: 0.003 and *T. brucei gambiense* DAL972:0.004) show less divergence than seen between *T. theileri* and *T. melophagium*.

**Figure 3:**
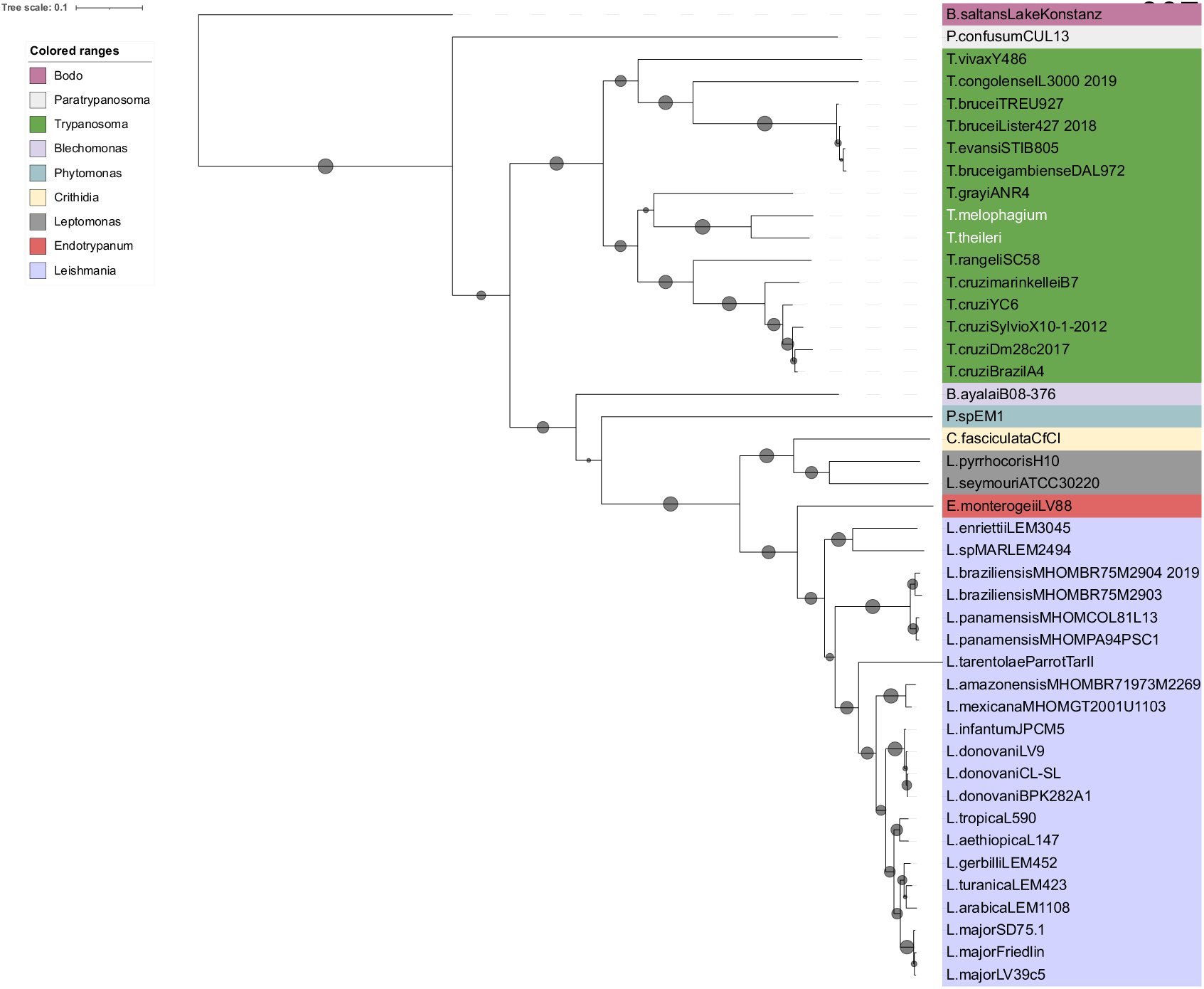
Species consensus tree based on 2,312 species trees created by STAG and STRIDE, OrthoFinder. The support values are represented by circles. Support values correlate to the proportion of times that the bipartition is seen in each of the individual trees used to create the consensus tree. The scale represents substitutions per site.

*T. theileri* has a greater number of species-specific orthogroups than *T. melophagium* and 12.9% of its genes are assigned to one of these orthogroups, while *T. melophagium* has only 2.7% of its genes in a species specific orthogroup (Table 2). Terminal branch length is correlated with specific orthogroup counts, which could account for the discrepancy. However, *T. melophagium* and *T. theileri* are each other’s most recent common ancestor and have been evolving at a roughly similar rate since this time, with similar terminal branch lengths (Fig. 3). To visualise these differences, the number of genes in each orthogroup was compared between *T. melophagium* and *T. theileri*. Orthogroups associated with host interaction protein families were highlighted based on their identification as a putative cell surface protein by Kelly *et al*. 2017 (32). Many of the *T. theileri* species specific orthogroups expansions belong to a cell surface family (Fig. 4a).

**Figure 4:**
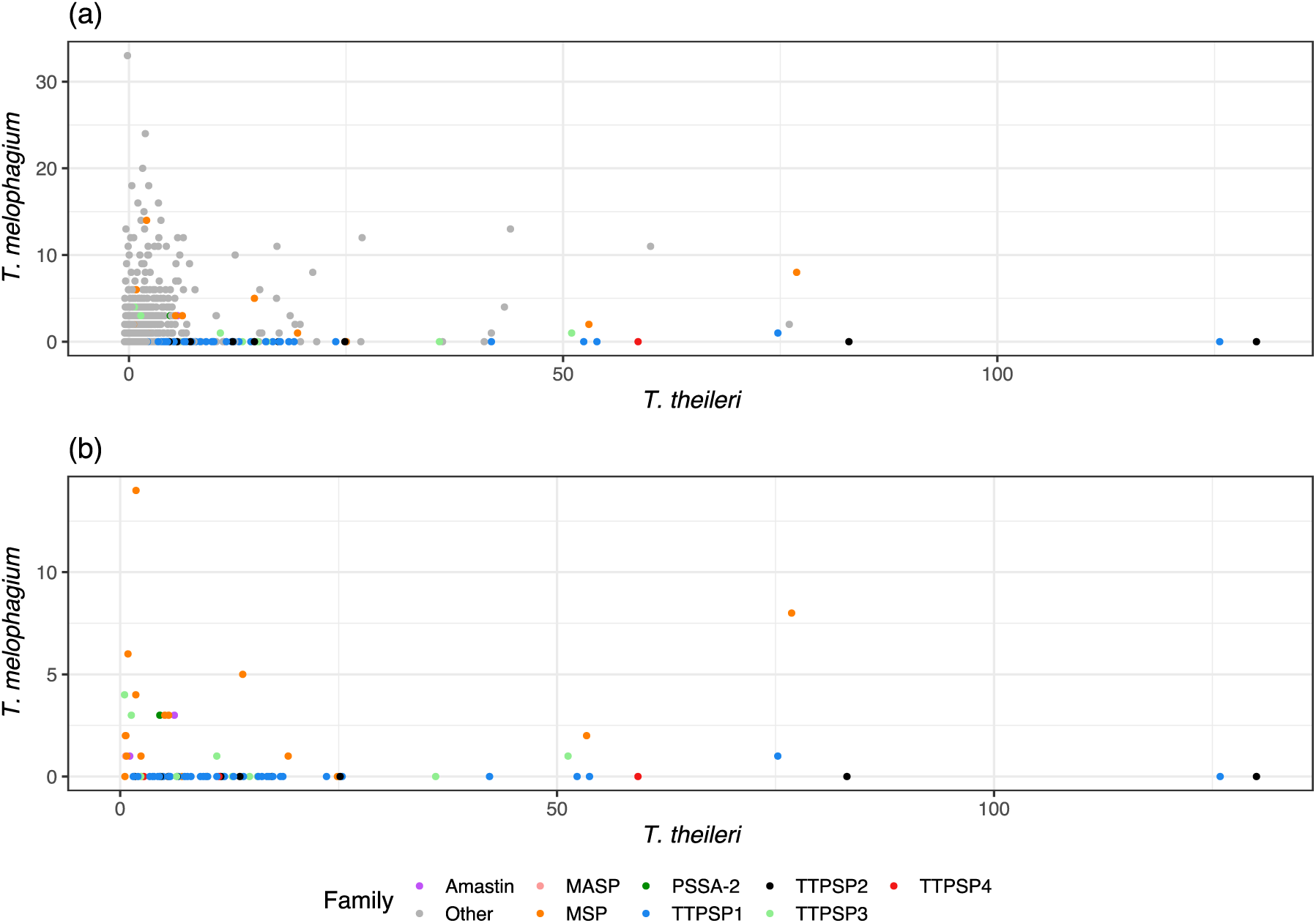
(a) Orthogroup and (b) host interaction orthogroup size comparison between *T. melophagium* and *T. theileri*. Each dot represents the numbers of genes found in each orthogroup for both species. The orthogroups have been annotated with their designation as either a putative cell surface protein family or ‘other’ (32).

**Table 2:**
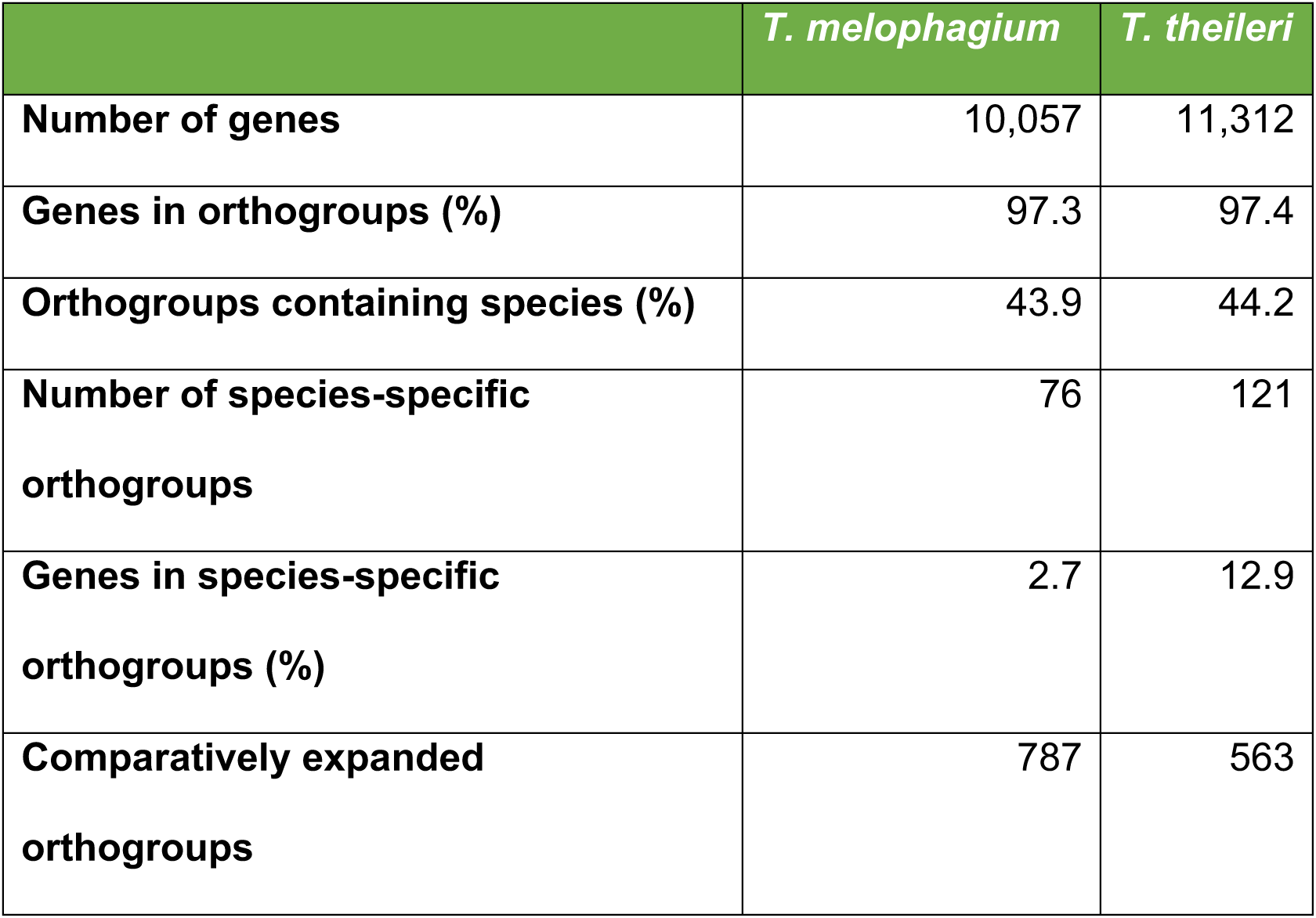
Orthologous protein clustering statistics of *T. melophagium* and *T. theileri*. The full species-specific clustering summary can be found in Supplementary File S3.

|  | *T. melophagium* | *T. theileri* |
| --- | --- | --- |
| **Number of genes** | 10,057 | 11,312 |
| **Genes in orthogroups (%)** | 97.3 | 97.4 |
| **Orthogroups containing species (%)** | 43.9 | 44.2 |
| **Number of species-specific orthogroups** | 76 | 121 |
| **Genes in species-specific orthogroups (%)** | 2.7 | 12.9 |
| **Comparatively expanded orthogroups** | 787 | 563 |

#### Interaction with the mammalian host and predicted cell surface proteins

Firstly, and as expected, *T. melophagium* was found to lack any genes in orthogroups which contained VSGs, characteristic of African trypanosomes (Supplementary File S4). To validate this, a blast search was performed using a relaxed cut off (1e-5) using the *T. melophagium* genome as the database and *T. brucei* TREU927/4 VSGs as the query. No hits were identified.

To enable comparison to the *T. theileri* genome analysis, the genes and orthogroups from this study were annotated with orthogroups from Kelly *et al*. (2017) (32). Across the entire genome, *T. theileri* is predicted to contain 1,265 more genes than *T. melophagium* (Table 1). Examination of orthogroups which were associated with a *T. theileri* putative surface protein (TTPSP) revealed a large expansion in *T. theileri* (1,251 genes) compared to *T. melophagium* (10 genes) which could equate to much of the disparity in genome size (Fig. 4). To confirm the difference in TTPSPs, the *T. theileri* transcripts were subjected to a blastn search against a database consisting of the *T. melophagium* transcripts (1e-25 cut-off). The *T. melophagium* transcripts were derived from the genome annotation analysis. Only 9 *T. theileri* TTPSPs aligned to *T. melophagium*.

TTPSPs were split between four orthogroups in the original *T. theileri* genome analysis but were split between 66 orthogroups in this analysis (Table 3) (32). TTPSPs share conserved C terminal GPI addition and N terminal signal sequences and contain regions of high divergence in the remainder of the sequence. TTPSPs are highly expressed as a family at the population level and are largely contained within tandem arrays, highlighting a similarity to the VSGs of *T. brucei* (32).

**Table 3:** Cell surface orthogroup counts from Kelly *et al*. (2017) (32) and this study along with counts of genes present in each category.

| Cell surface category | Annotation | Orthogroup count<br>Kelly *et al* / this study | Gene count<br>*T. theileri* / *T. melophagium* |
| --- | --- | --- | --- |
| Conserved | Amastin | 1 / 2 | 7 / 4 |
| Conserved | MASP | 1 / 1 | 1 / 1 |
| Conserved | MSP | 1 / 18 | 229 / 52 |
| Conserved | PSSA-2 | 1 / 1 | 5 / 3 |
| Unique | TTPSP1 | 1 / 42 | 720 / 1 |
| Unique | TTPSP2 | 1 / 10 | 301 / 0 |
| Unique | TTPSP3 | 1 / 11 | 157 / 9 |
| Unique | TTPSP4 | 1 / 3 | 73 / 0 |

Excluding *T. melophagium*, TTPSPs are absent from all other kinetoplastid species analysed (Supplementary File S4) revealing their specific innovation in these related trypanosomatids.

Following the TTPSPs, the largest expanded gene family in *T. theileri* are the MSPs. This protein family belongs to the peptidase M8 family of metalloproteinases. MSPs are likely to have contrasting roles in different life stages but their best understood function is to bind and cleave members of the complement system. This role presumably allows for evasion of complement-mediated proteolysis and therefore assists survival in the insect and mammalian hosts (23). Orthogroups which annotated as MSPs were expanded in *T. melophagium*, such as OG0008865. *T. melophagium* also has a species specific orthogroup (OG0009903) consisting of 11 proteins and annotated as leishmanolysin, indicative of cell surface protein families. Combined, these results suggest that the surface protein environment of *T. melophagium* and *T. theileri* are distinct and that genes encoding these proteins account for most of the discrepancy in the genome sizes.

#### Clade and species specific orthogroups

*T. grayi*, *T. melophagium* and *T. theileri* contain 42 clade-specific orthogroups including the MSP orthogroups OG0000590 and OG0008095. Both orthogroups were expanded in *T. theileri* in comparison to *T. grayi* and *T. melophagium*. *T. melophagium* and *T. theileri* share 81 orthogroups specific to the two species. Twelve of the orthogroups are putative cell surface protein families, including five representatives from TTPSP and seven MSP orthogroups. Of the nine-remaining annotated orthogroups, OG0008096 is the largest and contains trans-sialidsases. The orthogroup is expanded in *T. theileri* (n=17) compared to *T. melophagium* (n=5).

*T. theileri* has 121 species specific orthogroups. Sixty-four of these were putative host interaction genes (32). Two annotated orthogroups remained, which included a leucine rich repeat family associated with protein binding and (OG0011868) and a calpain cysteine protease family (OG0018204). *T. melophagium* has 76 species specific orthogroups, 68 of these were unannotated. Leishmanolysin families were annotated in three of the remaining orthogroups (OG0009903, OG0013746 and OG0013747). This annotation is associated with cell adhesion, such as MSP families. Other annotated species-specific orthogroups consisted of an actin family (OG0018163) along with several families associated with protein binding, WD domain G-beta repeat (OG0018131) and a leucine rich repeat family (OG0018098).

#### Modifying enzymes

Trans-sialidases are differentially expanded in *T. theileri* and *T. melophagium*, with 38 and 8 genes, respectively. Of the four orthogroups containing trans-sialidase, two do not contain proteins from *T. melophagium* (OG0011818 and OG0015016). These two orthogroups contain proteins from *T. cruzi* and so are likely to have been lost by *T. melophagium* and *T. grayi.* In *T. cruzi*, trans-sialidases are involved in host immune evasion (22).

Invertases are typical of the cell surface of *Leishmania* and is thought to transform sucrose into hexose in the gut of the vector. The orthogroups (OG0000150 and OG0000409) that include 20 *T. theileri* invertase genes only have 3 members in *T. melophagium* and 3 members in *T. grayi*. This expansion might indicate an adaptation of *T. theileri* to its vector, the tabanid fly, which can feed on sugary flower nectar (87). In contrast, the *T. melophagium* vector, the sheep ked, exclusively feeds on mammalian blood. The orthogroups with *T. theileri* invertases contain only one gene from the plant parasite *Phytomonas* (88, 89), suggesting a different mechanism for sucrose metabolism in these parasites.

Other putative cell surface modifying molecules, such as UDP-galactose/UDP-N-acetylglucosamine transferases (OG0000001) were expanded in *T. theileri* (n=60) compared to *T. melophagium* (n=11).

#### Glycolysis

Transcriptome comparison between *T. brucei* and *T. theileri* revealed differences in abundance of glycosomal enzyme mRNAs. Particularly, pyruvate orthophosphate dikinase, phosphoenolpyruvate carboxykinase and malate dehydrogenase were found to be >10 fold more abundant in *T. theileri* than in *T. brucei* (32). When comparing glycolysis-related genes in *T. melophagium* and *T. theileri*, we found that *T. melophagium* has expanded the number of genes in three of the orthogroups associated with glycolytic enzymes. These included pyruvate orthophosphate dikinase (OG0000570), phosphoenolpyruvate carboxykinase (OG0000120) and malate dehydrogenase (OG0000332), whilst *T. melophagium* has a reduced numbers of genes in the fumarate reductase orthogroup (OG0000078). Orthogroups associated with peroxisome targeting were found to be in equal numbers (OG0004108 PEX5 and OG0003998 – PEX7). All other glycolysis-related genes included in the comparison are present in equal numbers in the two species. Therefore, it is likely that the glycolysis pathway is conserved in *T. melophagium*.

#### Life cycle

Extensive studies of *T. brucei* have identified genes which are associated with key stages of the *T. brucei* life cycle. These studies tracked genes associated with stumpy formation in the blood stream form (90–93) and regulators of metacyclogenesis (94). These genes were combined with a list of validated development associated genes such as the RNA binding proteins RBP6, RBP7, RBP10 and ZFP2 and ZFP3 along with developmental regulators NRK A, NRK B, RDK1, RDK2, MAPK2 and phosphatases such as PTP1 and PIP39 (95–102). Most of the orthogroups containing these genes were represented with a similar number of genes in each species, indicating the presence of an environmental sensing ability and developmental competence (Fig. 5). However, there were notable differences. There is an expansion in the orthogroups containing KRIPP14, which is a mitochondrial SSU component (90), in *T. theileri*. *T. melophagium* has expanded its orthogroups containing the kinases NRK (96, 100), NEK and (99) ADKF (90) along with a dual specificity phosphatase (DsPho) and protein phosphatases 1 (PP1) (90, 103). Both *T. theileri* and *T. melophagium* are missing metacaspase (MCA1) which is associated with the later stages of progression towards metacyclic forms in *T. brucei* (94) and Hyp12 which upregulates bound mRNAs during development based on tethering assays in *T. brucei* (90, 103). Puf11, an effector molecule required for kinetoplast repositioning in epimastigotes (94), is also absent in *T. melophagium*.

**Figure 5:**
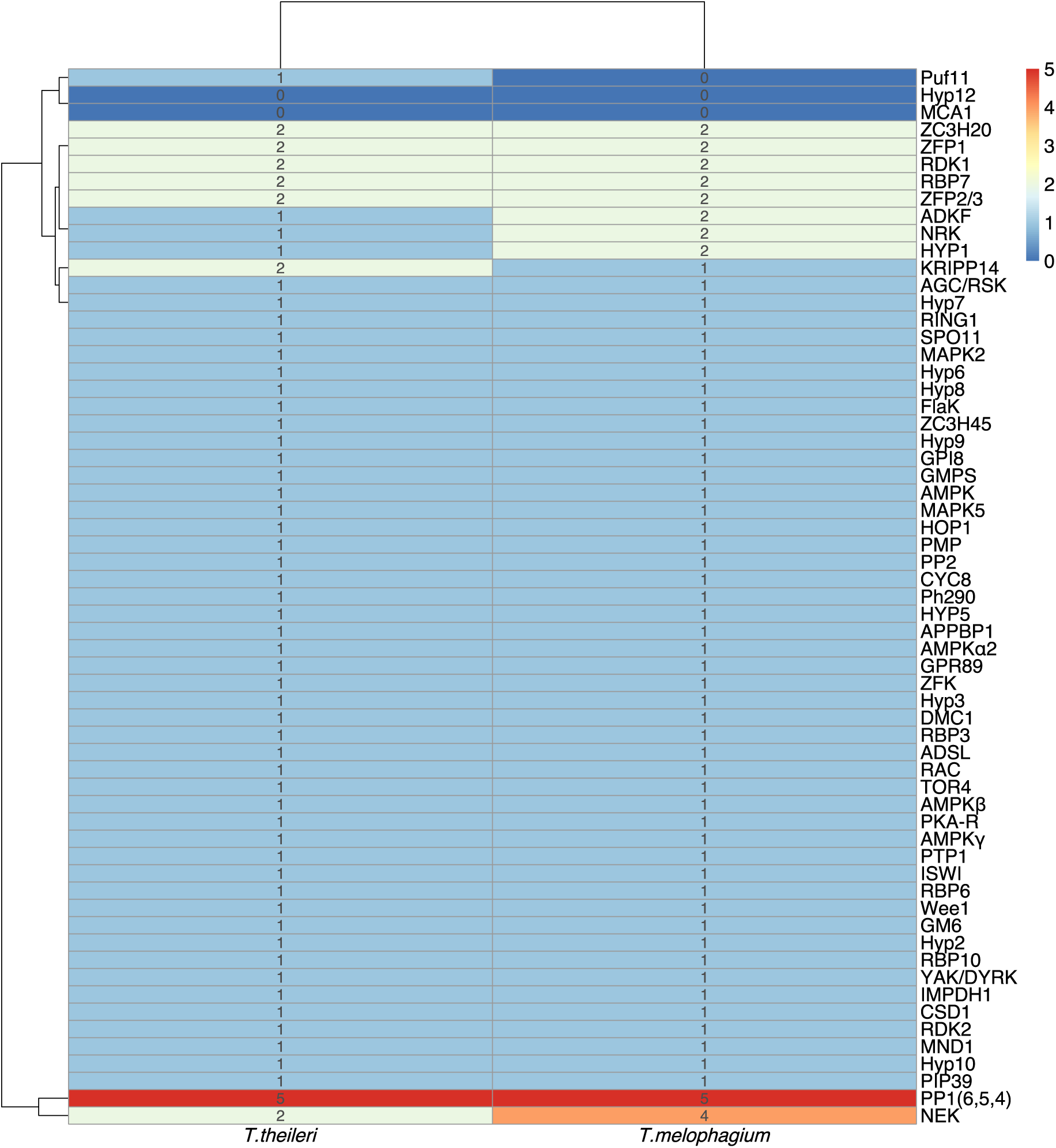
Genes associated with development, and related proteins, at various stages throughout the *T. brucei* life cycle. The number of genes in orthogroups associated with developmental regulation have been quantified in *T. theileri* and *T. melophagium*.

Life cycle regulatory genes and genes controlling meiosis SPO11, MND1, HOP1 and DMC1 were found in *T. melophagium* and *T. theileri* (Fig. 5), suggesting maintenance of a sexual stage.

#### RNAi

All 5 core genes that represent the trypanosome RNAi machinery (AGO1, DCL1, DCL2, RIF4 and RIF5) were present in *T. melophagium*, with an extra gene in the orthogroup containing DLC2 (Supplementary File S4). This indicates a functional gene silencing pathway is present in *T. melophagium* matching the prediction in *T. theileri*.

### 6.3. Discussion

*T. melophagium* and *T. theileri* are closely related trypanosomes that have distinct hosts and vectors (33, 34). Here, the genome of *T. melophagium* was sequenced and a draft assembly was produced and annotated. The annotated proteome was incorporated into a comparison with *T. theileri,* and other publicly available trypanosomatid proteomes, to determine their phylogenetic relationship and to explore the genomic basis of the host and vector specificity of these non-pathogenic trypanosomatids.

Although the two genomes compared in this analysis are predicted to be complete (Table 1), their assembly and annotation were performed five years apart using different pipelines and technologies. Comparing assemblies obtained using the same sequencing technologies and pipelines would allow for greater confidence in the observed variations in their genome content. This was not possible since many of the assembly tools used are specific for the sequencing technologies whilst there has been substantial development in assembly methods between the two studies. However, based on the convergence of the k-mer counting based prediction with the assembly sizes, along with 100% complete BUSCO scores, we were reassured by the quality of the draft assemblies and the subsequent comparisons of their genome content.

Using k-mer counting based predictions, *T. melophagium* was anticipated to have a smaller genome than *T. theileri*. This held true when the data were assembled (Table 1). The *T. melophagium* genome is more similar in size to *T. grayi*, the closest relative to *T. melophagium* and *T. theileri,* than to *T. theileri* (Fig. 2 & 3). *T. theileri* has likely expanded its genome size since speciation occurred. A peculiarity of *T. theileri* and *T. melophagium* is their highly reduced heterozygosity, in contrast to African trypanosomes (86). It is possible this reduced heterozygosity is linked to a founder effect (104). As *T. theileri* and *T. melophagium* have utilised specific host and vector niches, the small population that initially expanded into the niches possibly underwent a significant population bottleneck, especially as host domestication caused eradication of wild progenitors and wild relatives, which could have facilitated a reduction in heterozygosity, induced by genetic drift. Alternatively, the absence of a sexual cycle could contribute to the reduced heterozygosity. Although *T. melophagium* and *T. theileri* contain genes required for meiosis, it doesn’t confirm the species undergo sexual reproduction.

Selection appears to be acting to reduce the genome wide nucleotide biosynthesis cost in both *T. theileri* and *T. melophagium* (Fig. 2b) which has remodelled their genomes toward an AT bias, contrasting all other trypanosomatid genomes analysed in this study (Fig. 2 a & c). The predicted selection pressure acting to reduce nucleotide cost is at the expense of translational efficiency (Fig. 2c) and is greater than the free-living *Bodo saltans*, monoxenous insect parasites such as the early branching *Paratrypanosoma confusum* and *Phytomonas* EM1. *Phytomonas* has limited access to nitrogen as it infects nitrogen deficient plants and has been highlighted as an example where diet can cause selection to reduce the species genome nucleotide cost, through a reduction in GC content (76). We propose that the reduction in the selection cost of *T. theileri* and *T. melophagium* may be related to their non-pathogenic nature. By remodelling their genome to an AT bias, they may have reduced their cost to their host, facilitating reduced pathogenicity.

This clade specific genome remodelling provides an example of the similarity between *T. theileri* and *T. melophagium*. However, the species have contrasting hosts, vectors, and genome sizes. Genome annotation and orthology inference identified candidates for their discrepancy in genome size. When species specific orthogroups were compared, the greatest contrast was between orthogroups associated with the putative cell surface, with the largest expansions detected in *T. theileri* being of TTPSP and MSP surface protein families (Fig. 4). Although both species undergo a cyclical transmission cycle, which includes mammalian and insect stages, we hypothesise the respective prevalence in their mammalian hosts, and the contrasting life history of their respective vectors could explain the genome expansion in *T. theileri*. *T. theileri*, spread by tabanids, are found in over 80% of livestock (25–28). In comparison, *T. melophagium* exhibits lower detected prevalence, being rarely identified in its mammalian host via blood smears (34) or after blood culture (35). Moreover, sheep keds, which transmit *T. melophagium*, are intimately associated with their mammalian host, spending their entire life either on the sheep’s skin or wool. Here, males and females feed solely on mammalian blood providing many opportunities for transmission of *T. melophagium* from the sheep to the sheep ked (39). Therefore, there is potentially less advantage for *T. melophagium* to invest in mammalian immune evasion mechanisms required to extend the length of its infection in sheep, since it has many transmission opportunities. The limited investment in *T. melophagium* is emphasised by their relatively unsophisticated putative TTPSP repertoire alongside modestly expanded species-specific MSP families (Fig. 4). Instead, *T. melophagium* could rely on its ancestral ability to sustain infections in invertebrate hosts which, although able to be primed to defend against a specific pathogen, rely upon an innate immune response (105).

In contrast, *T. theileri* has a transient host-vector interaction. Tabanid flies of either gender survive on plant sugars whilst adult females occasionally feed on mammalian blood (42). Therefore, potentially *T. theileri* requires extended survival in its mammalian host to sustain transmission to cattle, compared to the intimate long-term association of sheep keds with *T. melophagium*. The investment from *T. theileri* in an expanded surface protein repertoire is likely to support adaptive immune evasion and prolonged survival in the mammalian stage of its life cycle. Alternatively, or additionally, differences between the bovine and ovine immune responses could contribute (106).

Many of the gene families identified in *T. theileri* (32) were further divided into multiple orthogroups in this study. The discrepancy is likely to be explained by evolution of the methods utilised by OrthoFinder. At the time of publication of the *T. theileri* study, OrthoFinder v.1 was available, while our analysis used version v.2.5. For this reason, we can speculate that the clustering in this study is more refined, such that the TTPSP families should be divided into smaller protein families. However, large paralogous orthogroups remain the toughest challenge for orthogroup clustering software and so the relationships between this set of proteins will continue to evolve alongside the software (107).

Genes involved in the trypanosome life cycle, cellular quiescence and meiosis were all detected in *T. melophagium*, suggesting a competent developmental cycle along with the machinery for sexual recombination. There is an expansion of the *T. theileri* invertase orthogroup which was not present in *T. melophagium*. This is potentially associated with the utilisation of sucrose in the tabanid fly’s diet. Although the glycolysis pathway is present in *T. melophagium*, there was an expansion in the pyruvate orthophosphate dikinase, phosphoenolpyruvate carboxykinase and malate dehydrogenase orthogroups. These genes are associated with the branch of the glycolysis pathway that converts pyruvate to succinate to facilitate the recovery of NAD^+^ (32). This branch of the glycolytic pathway was upregulated in *T. theileri* in contrast to *T. brucei* (32). Interestingly, the core RNAi genes were detected in *T. melophagium*, consistent with *T. theileri* but distinct from *T. cruzi* which is also a stercorarian trypanosome, but which lacks the requisite molecular machinery (108).

In summary, we have found that *T. theileri* and *T. melophagium* are closely related species which display substantial remodelling of their genomes to facilitate a reduction in their nucleotide costs, which could reduce the costs they impose on their hosts. *T. theileri* displays a considerable genome expansion which is associated with a large repertoire of unique proteins that characterise its cell surface and host interaction gene repertoire. These genes could facilitate a lifelong infection in its mammalian host. The comparatively unsophisticated immune evasion repertoire displayed by *T. melophagium* suggests that the species has emphasised its adaptation to the insect stage of its lifecycle.

## 7. Author statements

### 7.1. Authors and contributions

| Contributor Role | Author |
| --- | --- |
| **Conceptualisation** | GO, KM |
| **Methodology** | GO, BM |
| **Software** | GO |
| **Validation** | GO |
| **Formal Analysis** | GO, BM |
| **Investigation** | GO, BM |
| **Resources** | GO, KM, JLV |
| **Data Curation** | GO |
| **Writing – Original Draft**<br>**Preparation** | GO, BM |
| **Writing – Review and Editing** | GO, KM, BM, JLV |
| **Visualisation** | GO, KM, BM |
| **Supervision** | KM |
| **Project Administration** | GO, KM |
| **Funding** | KM |

### 7.4. Ethical approval

Not applicable.

### 7.5. Consent for publication

Not applicable.

## 7.6. Acknowledgments

We thank Steven Kelly (University of Oxford) for his comments and insights on the manuscript.

## 8. Supplementary material

**Supplementary Figure S1:**
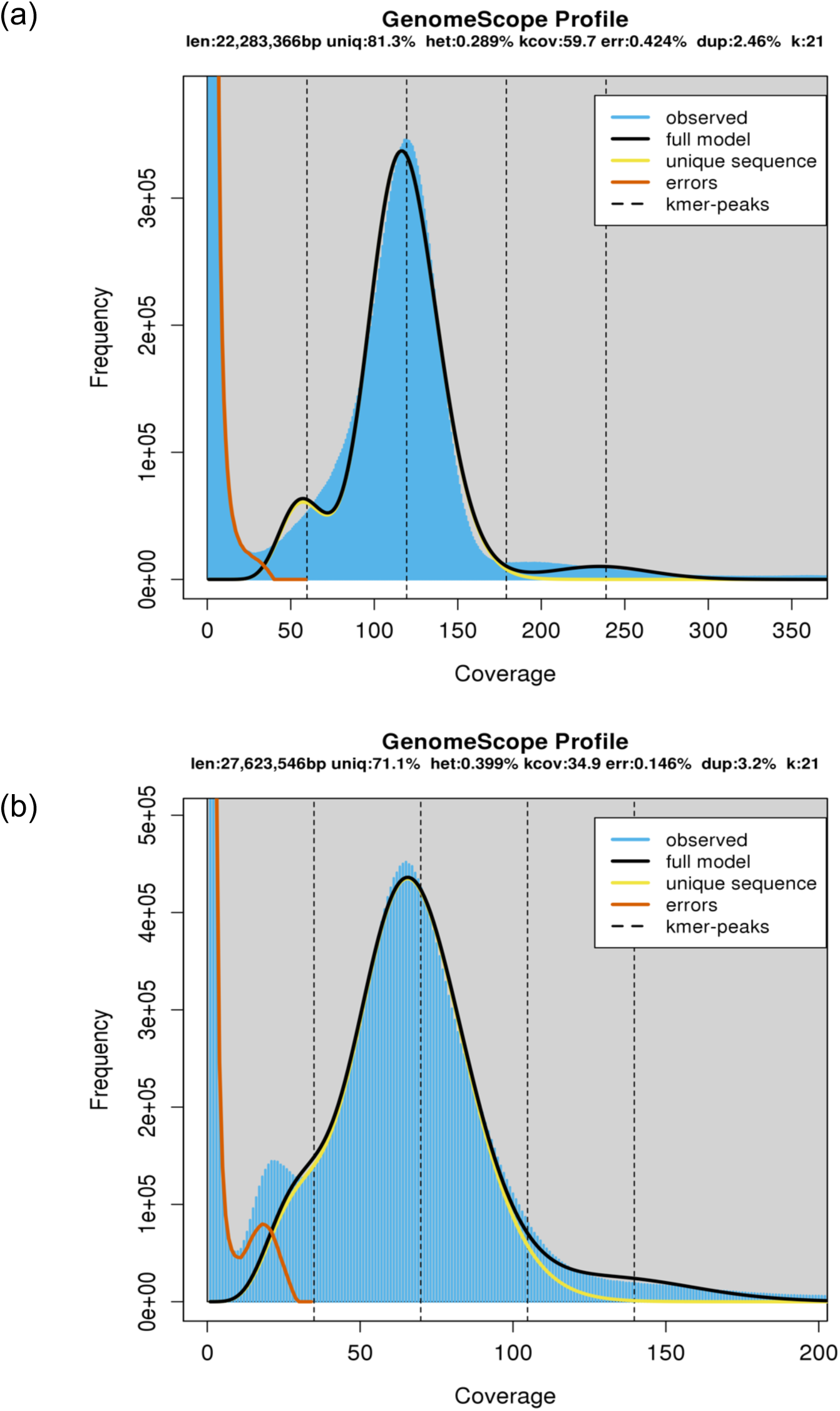
k-mer spectra of (a) *T. melophagium* and (b) *T. theileri* using a k-mer size of 21. The statistics presented in these images represent the lowest estimate size, the highest estimate was used in Table 1 as these had a greater model fit.

**Supplementary Figure S2:**
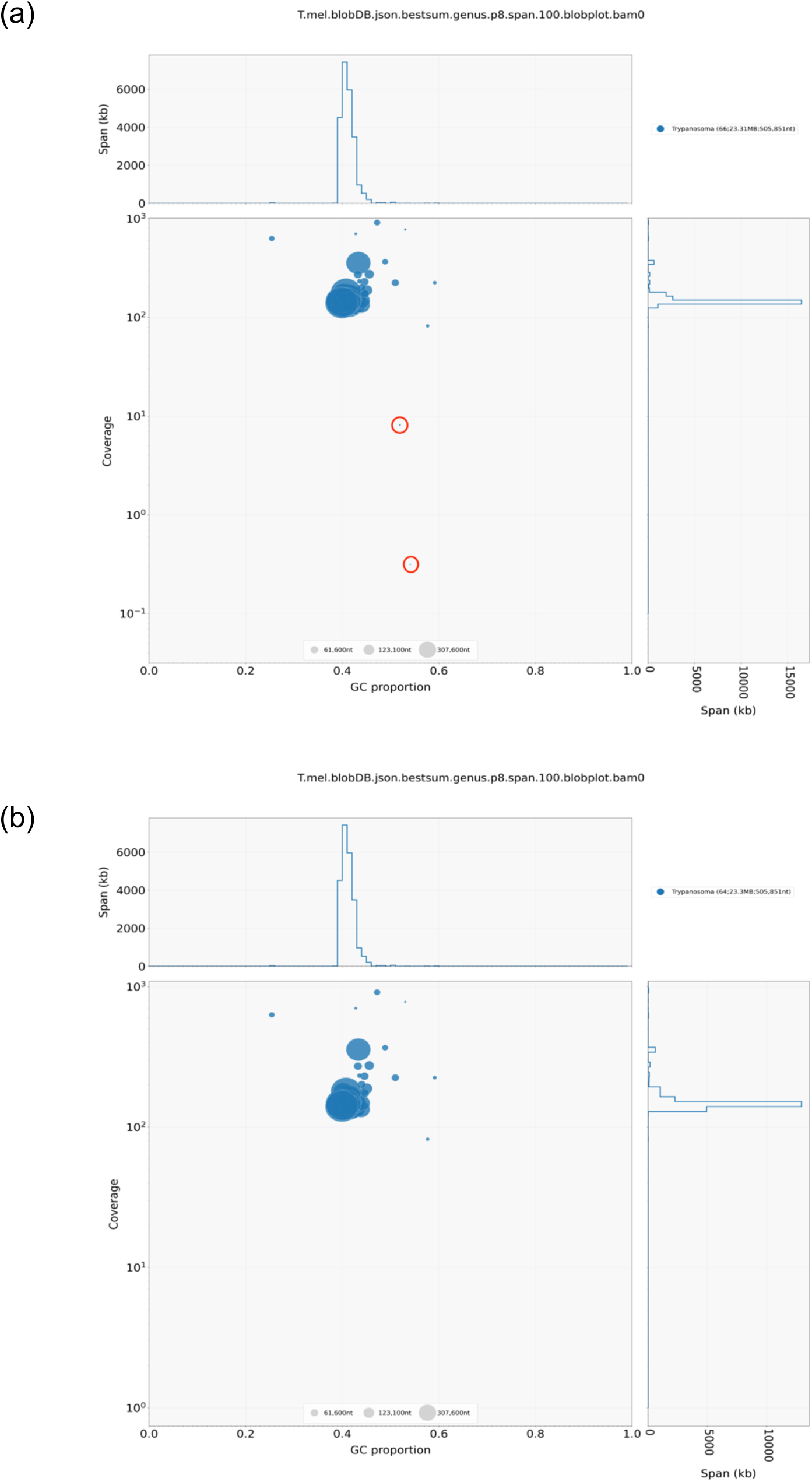
Coverage and blast annotation for (a) the polished assembly and (b) the manually trimmed assembly.

**Supplementary Figure S3:**
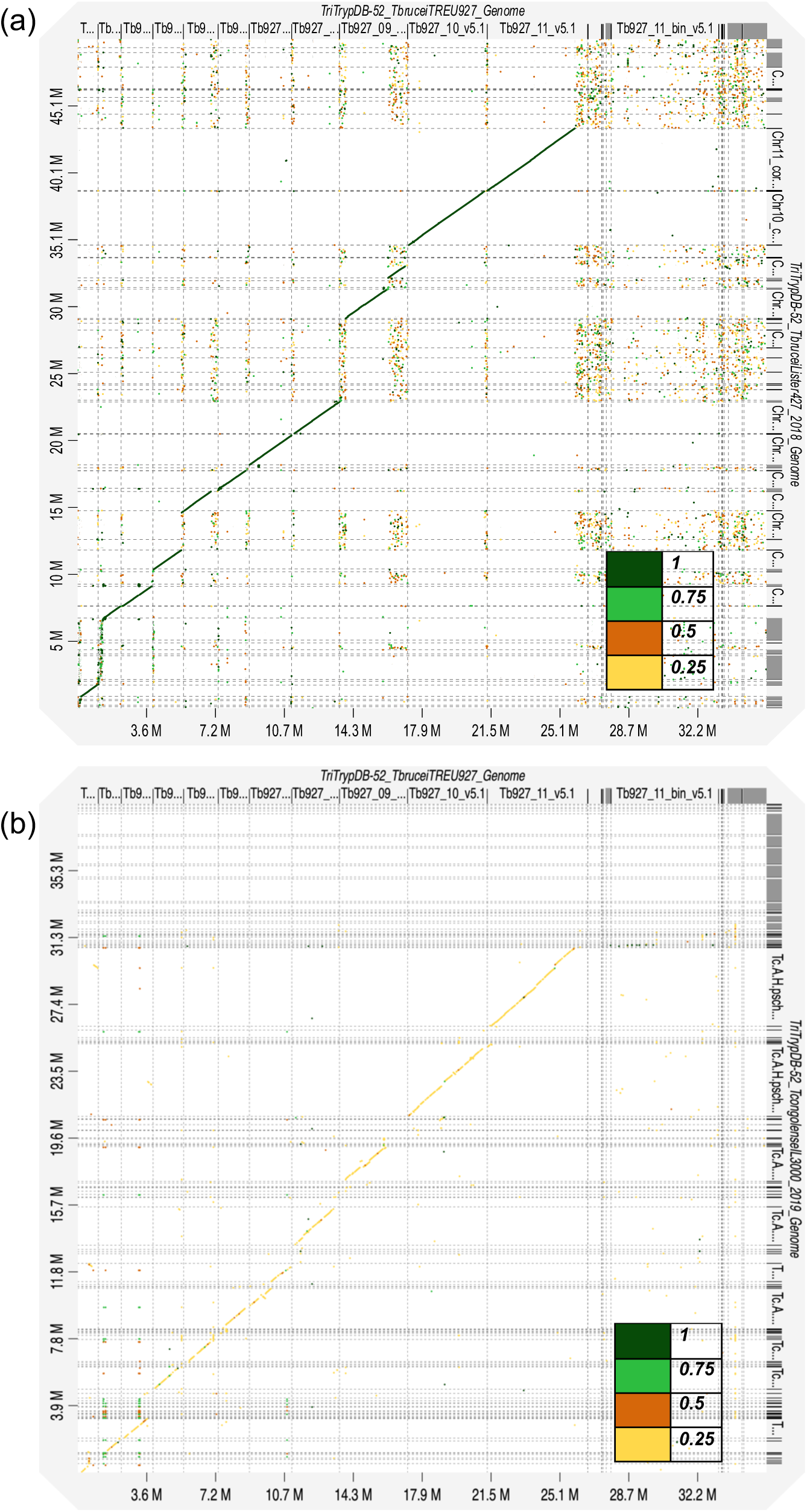
Synteny plot of (a) *T. brucei* TREU927/4 and *T. brucei* Lister 427 and (b) *T. brucei* TREU927/4 *and T. congolense* IL3000 2019 genome sequences. The legend refers to the identity between the sequences.

**Supplementary Figure S4:**
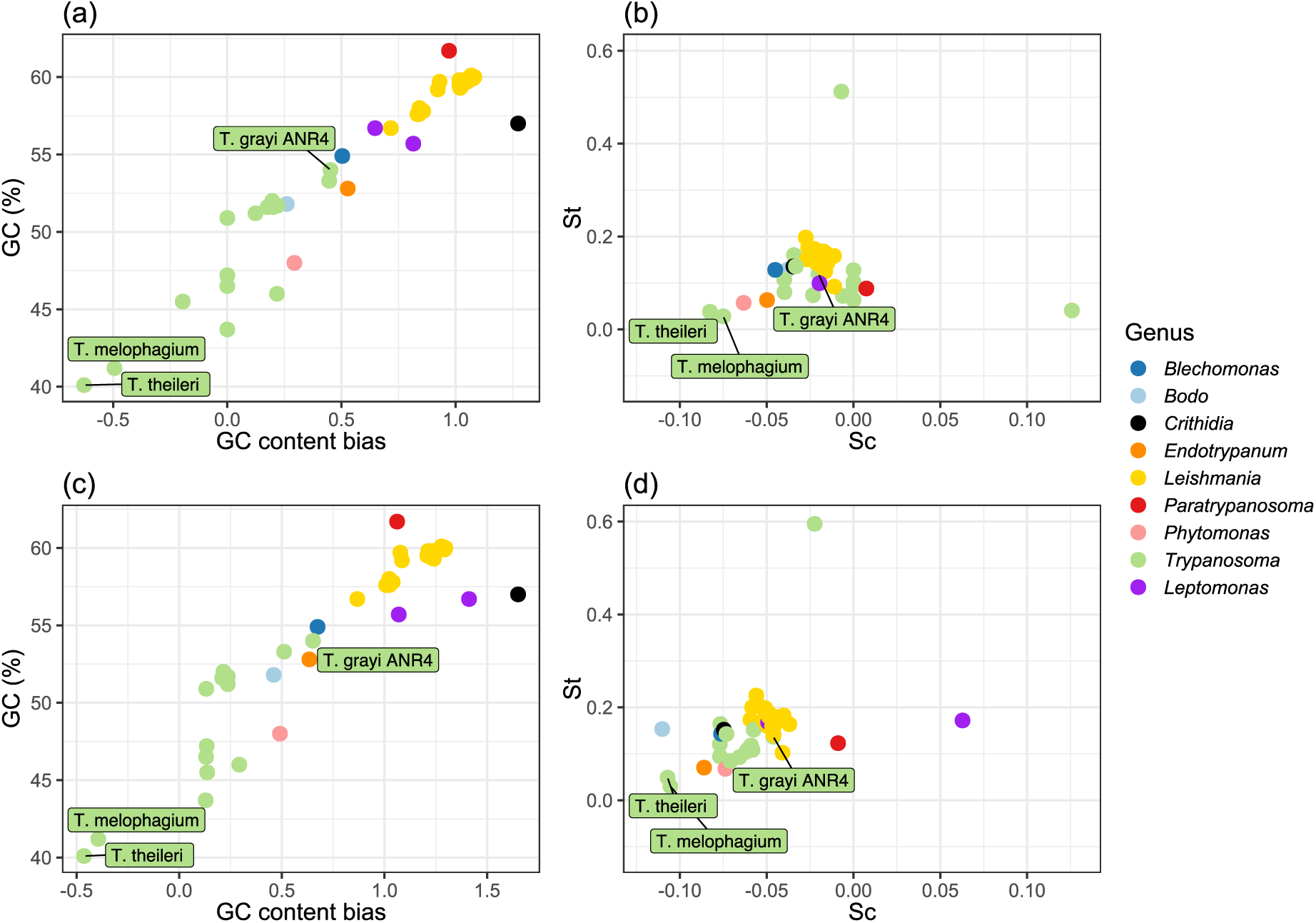
(a) GC content across the whole genome and GC content bias in a every CDS. (b) Selection acting on translational efficiency (St) and selection acting on nucleotide cost (Sc). (c) GC content across the whole genome and GC content bias in CDS of a trypanosomatid universal single copy orthologue which is essential in every life cycle stage in *T. brucei* (n=158). (d) Selection acting on translational efficiency (St) and selection acting on nucleotide cost (Sc) in trypanosomatid universal single copy orthologues which is essential in every life cycle stage in *T. brucei* (n=158).

**Table S1:**
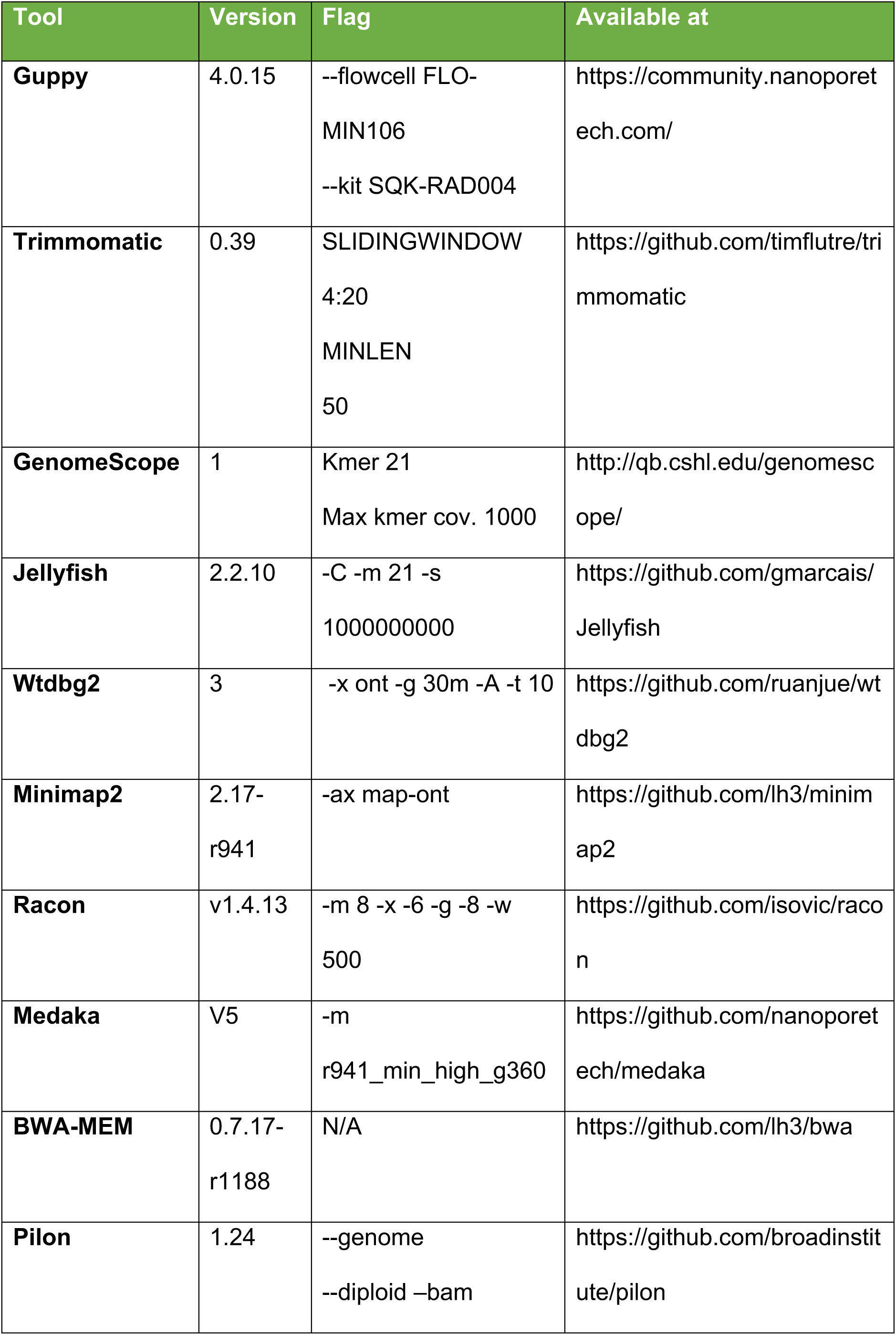
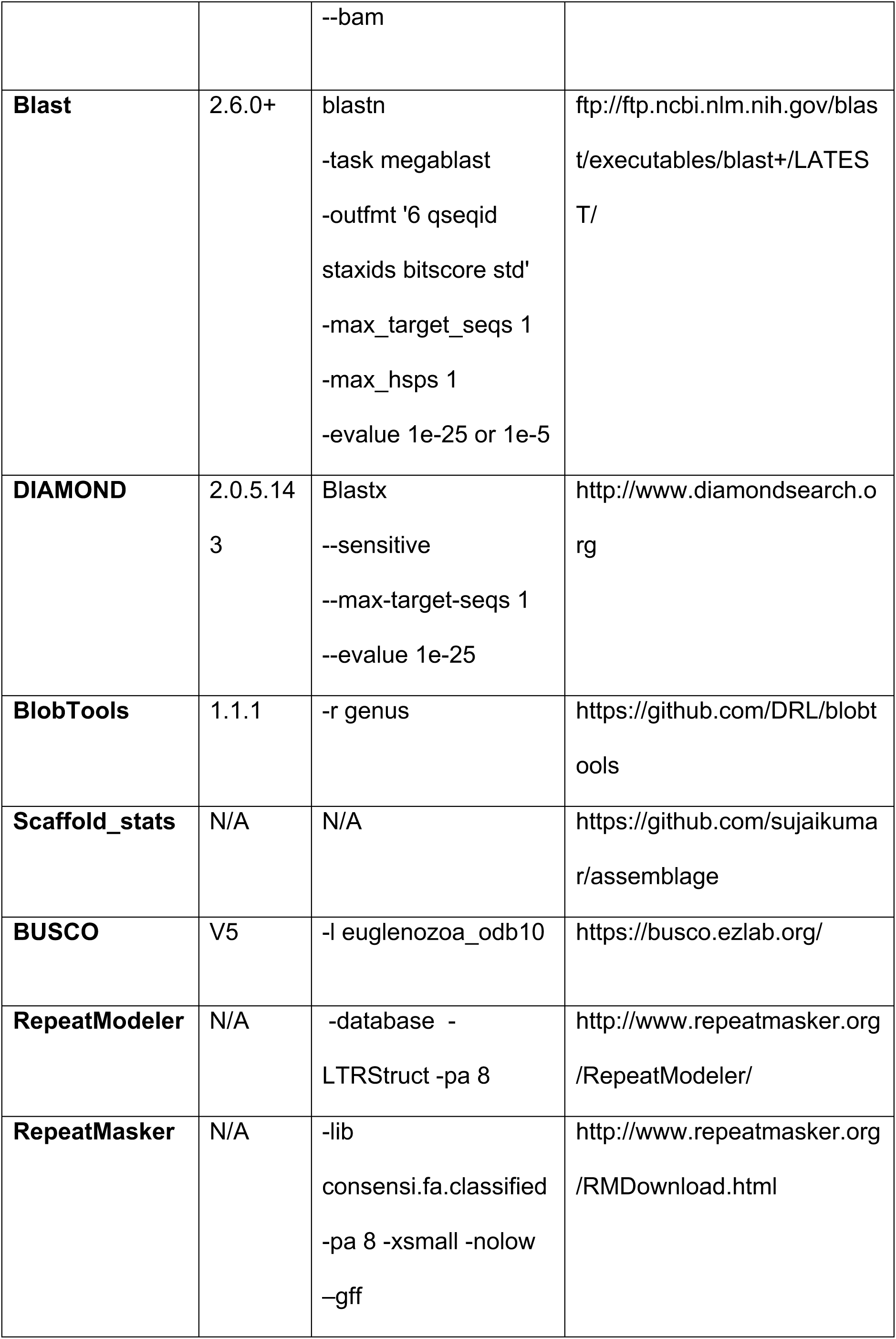
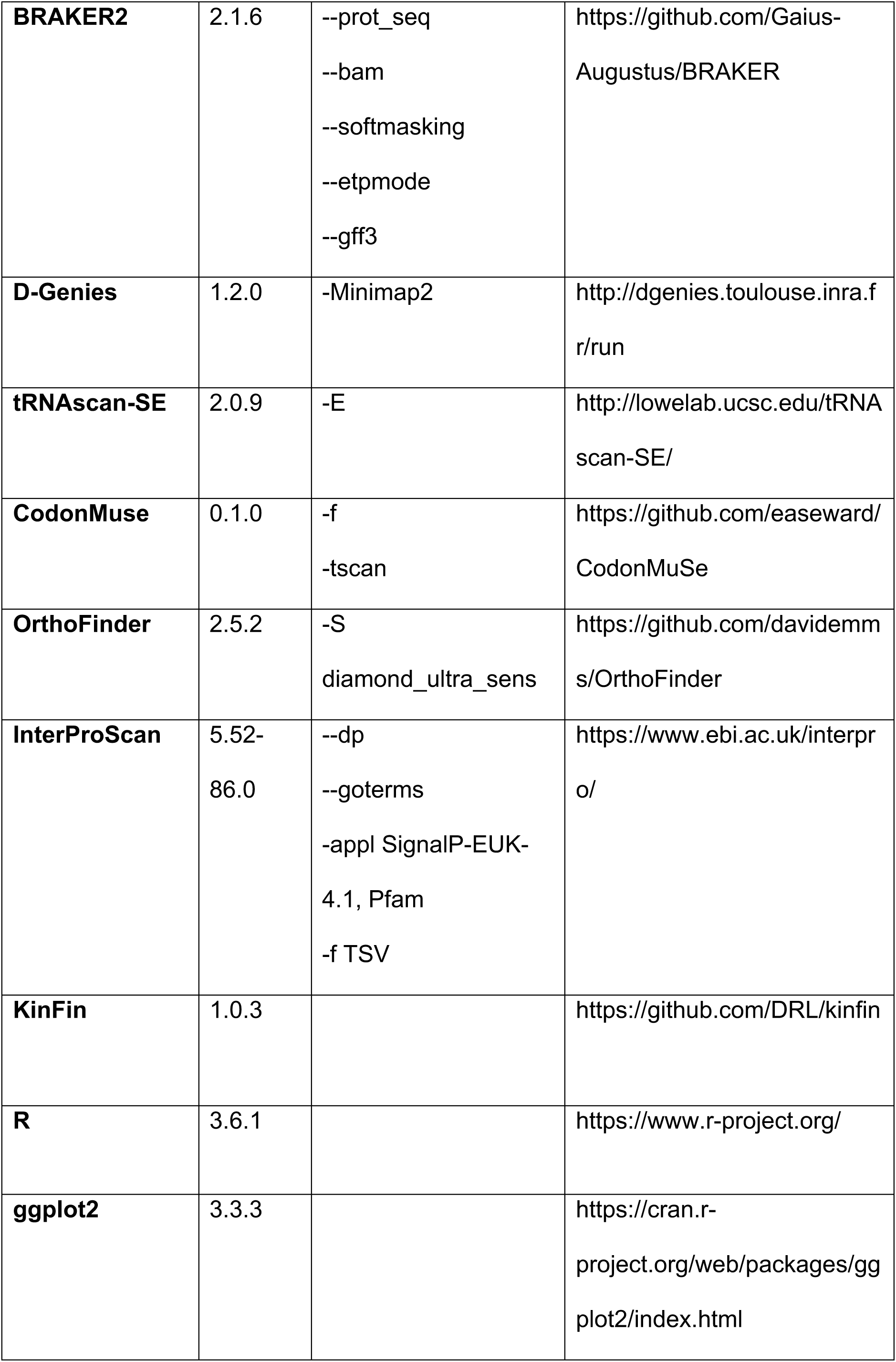
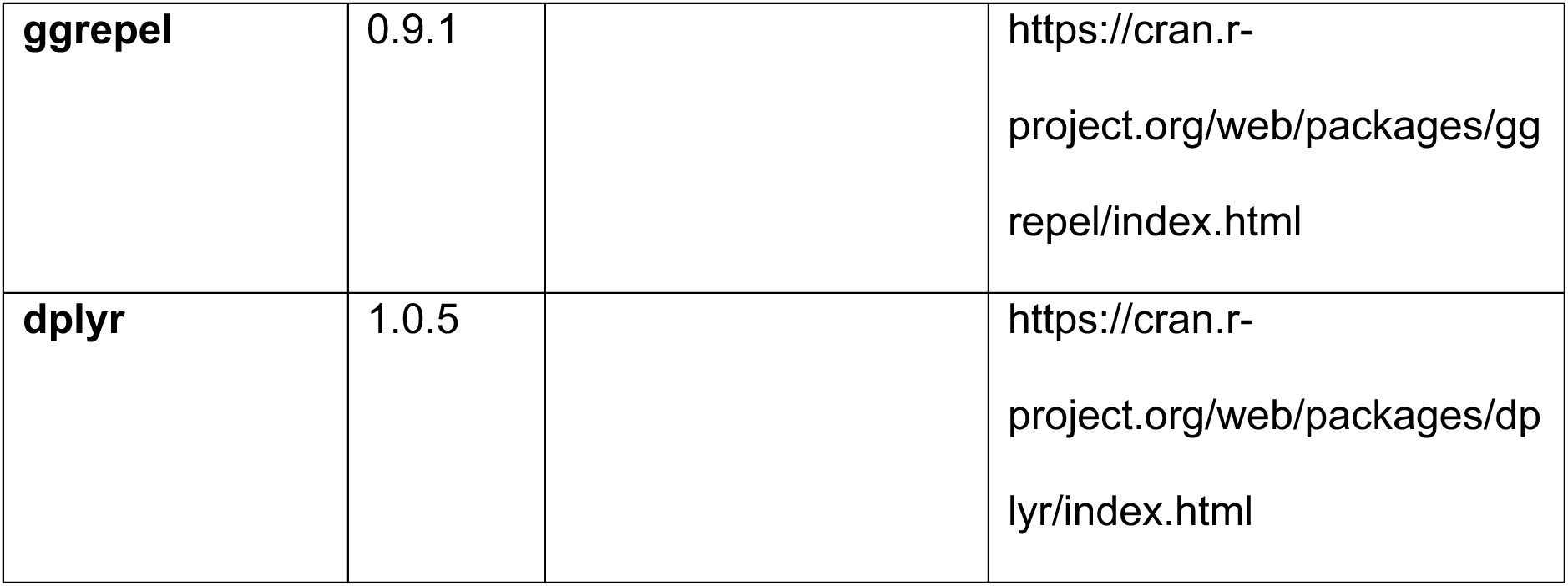
Software, including the version and any options used.

